# Trace elements increase replicability of microbial growth

**DOI:** 10.1101/2024.07.15.603609

**Authors:** Amit Shimoga Nadig, Rotem Gross, Tobias Bollenbach, Gerrit Ansmann

**Affiliations:** Institute for Biological Physics, University of Cologne, Zülpicher Straße 77, 50937 Köln, Germany; Center for Data and Simulation Science, University of Cologne, 50931 Köln, Germany

## Abstract

Trace elements are often omitted from chemically defined growth media. From established properties of trace elements, we deduce that this omission makes experiments unnecessarily sensitive to unavoidable contamination with trace elements. We confirm this experimentally by growing eleven bacterial strains in high replicate with and without supplementing trace elements, keeping all other conditions as fixed as possible to isolate the effect of trace elements. We find that supplementing trace elements considerably reduces variability of growth even in this benign scenario, and we argue that typical experimental setups exacerbate this. We discuss implications for the design and use of trace-element supplements, and in particular argue that their use should be standard practice, as they can reduce variability of almost all experiments using chemically defined media, taking a step towards greater precision and replicability in microbiology.

---

Reducing the variability of growth experiments is crucial in microbiology^1,2^, as it enhances the power of individual comparative studies and the comparability and replicability across studies. To this end, chemically defined growth media such as M9 or MOPS are widely used^3,4^. Low concentrations of trace elements in such media are beneficial for many microbes and for realism^3–5^. Historically, tap water was used to prepare the medium and provided trace elements, but when purified water is used for replicability, supplementing them is recommended^6,7^. However, many studies and protocols omit them^7–10^, sometimes deliberately^11–13^.

If trace elements were consistently absent, replicability would not suffer. However, a concentration of zero is an unattainable ideal. Possible sources of contamination include impurities in ingredients^13–15^ and labware^16^, production and cleaning residues on labware^17^, and aerosols^18^. Since trace elements already take effect at very low levels, we expect such contamination to cause considerable experimental variability (Fig. 1). Supplementing trace elements moves an experiment away from a point of high sensitivity and therefore reduces the relative impact of contamination. We therefore hypothesise that it generally increases the replicability of microbial growth.

**Fig. 1.**
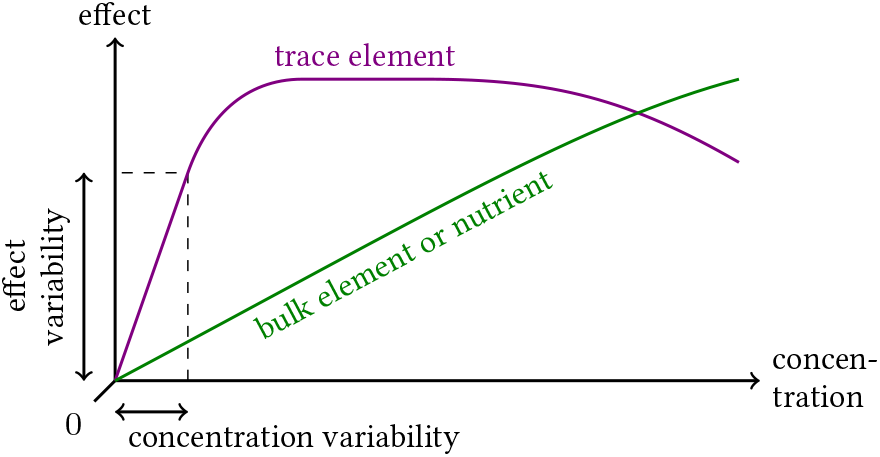
The impact of contamination can be reduced even when it is unavoidable or unknown. Schematic of the impact of concentration variabilities, which scales with the derivative of the effect with respect to concentration (akin to classical error propagation^19^). The impact is therefore highest for trace elements at zero concentration, and lower for bulk elements and nutrients as well as for trace elements at higher concentrations.

This is a systemic effect that relies only on the established property of trace elements to take effect at low concentrations. It therefore also covers adverse effects as well as microbes whose metabolism is only poorly understood so far. On the other hand, due to the stochastic nature of contamination and the diversity of underlying low-level mechanisms, it is not feasible to predict the occurrence and magnitude of this effect. Demonstrating it therefore requires considering statistical trends in a high number of dedicated experiments.

To test our hypothesis experimentally, we performed bacterial growth experiments in microtitre plates in high replication, assessing variability with and without trace-element supplementation (Fig. 2, left panels). We used a standard supplement^4^ containing manganese, iron, zinc, cobalt, boron, copper, aluminium, nickel, molybdenum, tungsten, and selenium (Ext. Tab. I). To avoid other, confounding sources of variability, we first inoculated a large volume from a pre-culture in exponential phase under the same conditions, which we then divided into the several replicates, incubated simultaneously in a homogenised environment, and measured with a robot. This approach minimised differences due to inoculum size, adaptation to new conditions (lag phase), well location, and human error (Methods 3).

**Fig. 2.**
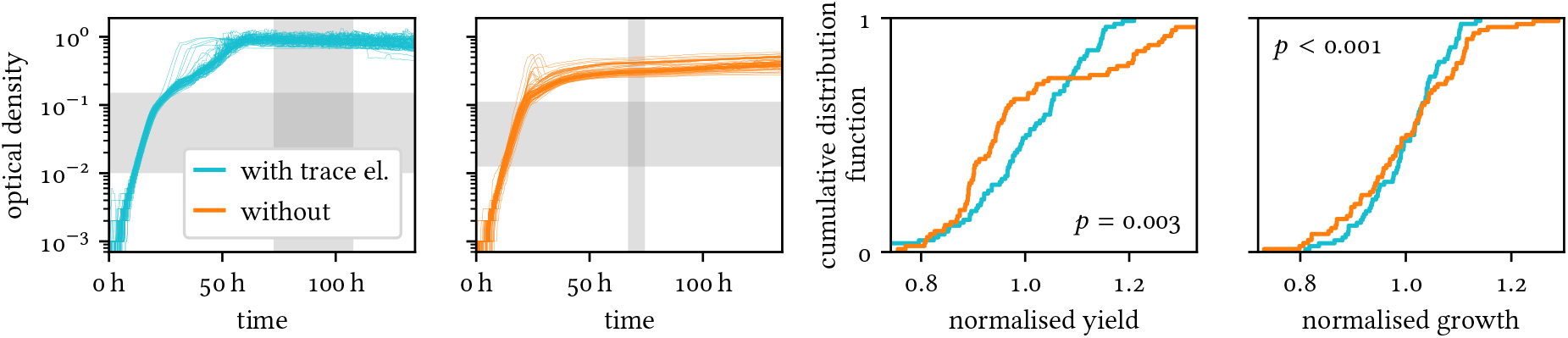
Trace elements reduce growth variability in *Sphingomonas molluscorum*. Left panels: 80 growth curves of a *Sphingomonas molluscorum* strain recorded under identical conditions with and without supplemented trace elements. Grey areas mark the automatically chosen ranges of time and optical density used to determine yield and growth rate, respectively (Methods 6). Curves are smoothed with a length-5 median filter (unsmoothed curves in Ext. Fig. 2). Right panels: Distributions of normalised growth characteristics with and without trace elements with the one-sided *p* value for the null hypothesis that trace elements have no effect on the coefficient of variation (Methods 7).

We used the coefficient of variation to estimate relative variability of growth rate and yield. Following common practice, we determined growth rate from a fit in an interval of optical density (as a proxy for biomass)^20^. To avoid bias in the selection of this interval, we automated it with the objective of a high goodness of fit and a low variability of the resulting growth rates (Methods 6). Similarly, we determined yield from a time interval that was automatically chosen with the objective of stagnant growth and low variability.

To avoid capturing any species-specific peculiarities, we repeated this procedure for eleven strains from different species, representing both Gram-negative and -positive bacteria (Ext. Tab. II, Ext. Fig. 1, and Methods 2). We verified that there is no significant correlation between the absolute measures (growth and yield) and their variabilities (Ext. Fig. 4). This excludes certain artefacts, e.g., trace elements leading to higher yields, which in turn can be measured more accurately.

Trace elements reduced the variability of yield and growth rate. For the example of *Sphingomonas molluscorum* (Fig. 2), yield variability dropped from 16% to 12% and growth-rate variability from 11% to 7%. Across all strains, the median variability of yield and growth rate was reduced by about a third each, with the effect being significant for six strains for yield, nine strains for growth rate, and overall for both measures (Fig. 3). The medians of variabilities over both conditions were 1.7% for yield and 7% for the growth rate. These rather low values most likely reflect our efforts to minimise other, confounding sources of variability and that we optimised the determination of yield and growth rate for low variability. Under more typical experimental conditions, we expect that more contaminations and interacting causes will increase the absolute variability difference, but also make it more difficult to disentangle it from other influences. We conclude that supplementing trace elements generally reduces the variability of microbial growth.

**Fig. 3.**
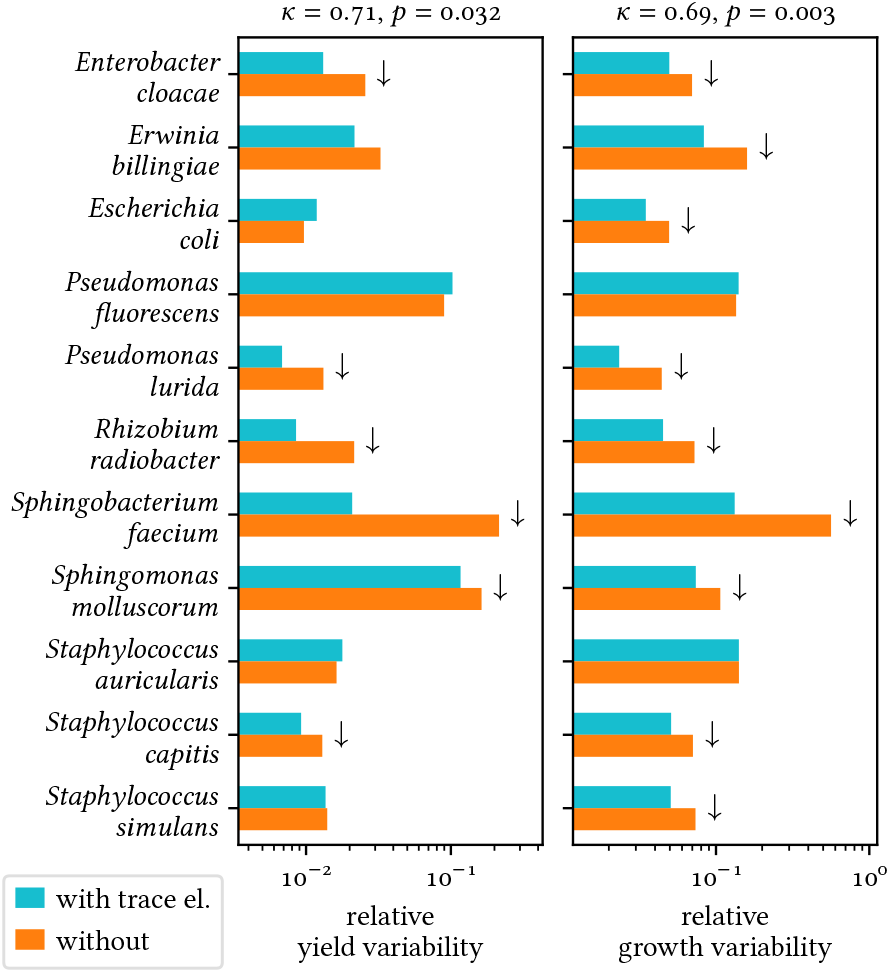
Trace elements reduce growth variability for many strains. Coefficients of variation of yield and growth rate with and without trace elements. Arrows indicate significant differences for individual strains (α = 0.05, two-sided). κ is the median ratio (with by without) of the respective coefficients of variation, and the *p* value is for the null hypothesis that κ = 1 (two-sided Wilcoxon’s signed-rank test, Methods 7).

According to our hypothesis, the increased variability without supplemented trace elements is caused by differences in contamination between the wells. We excluded contamination to the glassware that withstood mixing (Ext. Fig. 3). Instead, we showed that contamination originates either from the microtitre plates or the experimental procedure itself, possibly from aerosols: We applied our incubation and measurement procedure to highly purified water instead of inoculated medium and then used mass spectroscopy to compare its elemental content with that of negative controls that did not undergo the procedure (Methods 8). We found that the procedure increased the concentration of the supplemented trace elements overall, in particular for boron (4.5× or more) and manganese (2.4×) (Ext. Fig. 5 left). Notably, for those elements featured in the supplement we used, the supplemented concentrations were higher. This observation aligns with our hypothesis that supplementing trace elements mitigates the impact of contamination (Fig. 1).

Although we kept the conditions in our experiment as fixed as possible, supplementing trace elements already had a noticeable effect, e.g., reducing median variabilities by one third. While this effect was small or absent for some strains, this is unlikely to hold for all conditions. For instance, when a microbe requires a trace element to respond to a particular stress, contamination of that element may become important. Also, even small absolute increases in variability may suffice to mask a large effect in combination with other confounding factors or a small effect that is indicative of something larger, e.g., in an exploratory screening. Moreover, this issue is likely to become more relevant when differences in ingredients, labware, equipment, or the lab environment cannot be avoided, e.g., when comparing studies or in a long-term experiment^12^. This is corroborated by recent observations of contaminations in sodium phosphate, particularly of iron and nickel, and that these affect the growth of *Escherichia coli*^15^. A standard trace-element supplement adds iron and nickel at much higher concentrations and would therefore mostly mitigate this problem. In another example, the *E. coli* Long-Term Evolution Experiment relies on iron being supplied by the water used for the medium, and changes in water quality are a concern and may have caused one extinction^12^. In hindsight, supplementing iron and possibly other trace elements would have resulted in an environment that is more stable– not in atom numbers but in terms of biological impact.

So far we assumed contaminations that equiprobably affect all parts of an experiment. However, a worst case worth considering is that they are tied to an investigated condition, e.g., when studying the effects of a drug that is contaminated. Supplementing trace elements mostly prevents systematically false findings arising in such a scenario. Similarly, we found that the effect of contaminations on some strains may be far above the typical one (e.g., *Sphingobacterium faecium* in Ext. Fig. 2). These considerations and results suggest that the distribution of effect sizes has a heavy tail and that it is ill-advised to only consider the typical effect when deciding whether to supplement trace elements, as your mileage may vary considerably.

Trace-element supplements are inexpensive and easy to procure. By contrast, it is often impractical and unreliable to minimise contamination by identifying its sources through elemental analysis (as in Ref. 15). Likewise, assuming that contamination is negligible would necessitate a study such as the one conducted here – for all relevant conditions. While supplements occasionally have unexpected negative effects on certain microbes^21,22^, a positive effect should be at least as likely and may be a welcome bonus. Furthermore, trace elements increase the realism of growth conditions as microbes have access to them in most natural environments. We therefore argue that adding trace elements to chemically defined media should be standard practice in microbiology (and beyond) and only be omitted in justified exceptions^13^.

Common supplements are designed to provide essential or beneficial elements for a wide range of microbes^4^. However, other elements can contaminate experiments as well (Ref. 15 and Ext. Fig. 5, right), and most of them affect at least some microbes^23,24^. We therefore advocate the development of improved supplements that include relevant contaminants – whether elements or other biologically important substances – at concentrations that compromise between improving replicability and harming microbes.

## Methods

*Experiment* refers to all growth experiments of the same strain under the same conditions, i.e. medium.

**Ext. Tab. I.** Trace elements (top) and other components (bottom) as contained in the supplement (ATCC, MD-TMS; also see Ref. 4), of which we added 1% to those media with trace elements.

| component | concentration |  |
| --- | --- | --- |
| MnSO<sub>4</sub> · H<sub>2</sub>O | 0.5 g/l | 3.0 mM |
| FeSO<sub>4</sub> · 7 H<sub>2</sub>O | 0.1 g/l | 0.36 mM |
| ZnSO<sub>4</sub> · 7 H<sub>2</sub>O | 0.1 g/l | 0.35 mM |
| Co(NO<sub>3</sub>)<sub>2</sub> · 6 H<sub>2</sub>O | 0.1 g/l | 0.34 mM |
| B(OH)<sub>3</sub> | 0.01 g/l | 0.16 mM |
| NiCl<sub>2</sub> · 6 H<sub>2</sub>O | 0.02 g/l | 84 µM |
| Na<sub>2</sub>MoO<sub>4</sub> · 2 H<sub>2</sub>O | 0.01 g/l | 41 µM |
| CuSO<sub>4</sub> · 5 H<sub>2</sub>O | 0.01 g/l | 40 µM |
| AlK(SO<sub>4</sub>)<sub>2</sub> | 0.01 g/l | 39 µM |
| Na<sub>2</sub>WO<sub>4</sub> · 2 H<sub>2</sub>O | 0.01 g/l | 30 µM |
| Na<sub>2</sub>SeO<sub>3</sub> | 0.001 g/l | 5.8 µM |
| NaCl | 1 g/l | 17 mM |
| MgSO<sub>4</sub> · 7 H<sub>2</sub>O | 3 g/l | 12 mM |
| CaCl<sub>2</sub> | 0.1 g/l | 0.90 mM |
| EDTA | 0.5 g/l | 1.7 mM |

### 1. Growth medium

Our medium was composed of M9 salts (Sigma M6030) supplemented with 2 mM MgSO_4_ (Honeywell M7506) and 0.1 mM CaCl_2_ (Honeywell 223506). For this study, we do not categorise magnesium and calcium as trace elements, although our general argument also applies to them. We used 4 ^g/l^ glucose (Sigma G8270) as a carbon source. The above components were prepared in higher concentration separately and autoclaved, except glucose, which was sterile-filtered. For all experiments with trace elements, we added 1% trace-element supplement (ATCC, MD-TMS; Ext. Tab. I). Otherwise, we added the same amount of water.

### 2. Bacterial strains

We selected twelve strains considering availability, phyletic distance (to avoid pseudo-replication), biosafety level, and whether they grow in our medium (Ext. Tab. II).

### 3. Growth conditions and measurements

For each strain and medium (with and without trace elements), we grew a pre-culture under the same condition as for the main measurement (see below) and inoculated from its exponential phase (determined in a pre-experiment) to reduce the need for adaptation and the additional variability arising from it. We inoculated a bottle of the respective growth medium with a ratio of 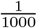, mixed it well, and transferred the result to a reservoir (VWR 732-1390) and from there to 96-well microtitre plates (Thermo 260860) with a pipetting robot (HighRes 1071562 with HighRes 1071667 tips), using 200 µl per well. We treated both experiments for a given strain as equally as possible, in particular we used the same batch of sterilised medium (except for adding trace elements or water as a last step) and allocated the experiments in equal parts (in different columns) onto each plate to reduce the impact of any plate differences. The plates were grown in a shaking incubator (Liconic STX44-HRSA) set to 30°C and 97% humidity (to minimise evaporation) and their optical density (OD) at 600 nm was measured every twenty minutes with a spectrophotometer (Biotek Synergy H1-MFDG) without lid, all by a robotic system (HighRes MC642).

**Ext. Tab. II.** Strains used in the main experiments.

| code | species | reference |
| --- | --- | --- |
| CEent 1 | *Enterobacter cloacae* | 25 |
| MYb 121 | *Erwinia billingiae* | 25 |
| MG 1655 | *Escherichia coli* | 26 |
| MYb 49 | *Ochrobactrum anthropi* | 25 |
| UTI 5 | *Pseudomonas fluorescens* | 27 |
| MYb 11 | *Pseudomonas lurida* | 25 |
| UTI 29 | *Rhizobium radiobacter* | 27 |
| MYb 181 | *Sphingobacterium faecium* | 25 |
| JUb 134 | *Sphingomonas molluscorum* | 25 |
| UTI 57 | *Staphylococcus aureus* | 27 |
| UTI 74 | *Staphylococcus capitis* | 27 |
| UTI 45 | *Staphylococcus simulans* | 27 |

In an alternative setup (Ext. Fig. 3), we reduced the amount of medium in the above preparation to only yield 100 µl per well. The other 100 µl came from one of eight bottles, into which we distributed the remaining medium (via VWR 732-1392). We used each of those eight bottles in equal parts for each plate, strain, and condition. We performed these experiments separately from the previous ones.

### 4. Preparing growth curves

We excluded Strain MYb49 *(Ochrobactrum anthropi)* from further analysis since – both, with and without trace elements – it grew badly and was subject to high measurement noise and extreme fluctuations (by an order of magnitude), such that it would be unsuitable for most quantitative analyses under these conditions.

For each experiment, we determined the baseline OD, i.e., the expected OD of the medium, as the median of the minima of each growth curve. We then subtracted this baseline OD from all measured ODs within the same experiment. For each growth curve, we removed all time points before the first local minimum, as those do not capture the growth dynamics of interest and are instead likely to be caused by initial condensation.

We excluded certain wells by manually inspecting two kinds of candidates: First, we considered wells whose minimal OD was larger than 0.01, indicating that they were subject to initial condensation. Second, to exclude wells that were likely biologically contaminated, we first determined a preliminary yield of each growth curve as the 98th percentile of its values and then considered growth curves whose preliminary yield exceeded the 90th percentile of all preliminary yields in the experiment by more than 50 %. Of 25 candidates identified this way, we excluded 22 (black in Ext. Figs. 1 and 2).

### 5. Quantifying variability

We quantified variability via the coefficient of variation, 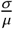, where σ is the standard deviation and μ is the mean of the respective measure. We opted against a ranked statistic since we expect trace-element contamination to cause meaningful outliers.

### 6. Growth characteristics

We computed the yield of a given growth curve as the 98th percentile of values within a given interval (vertical grey area in Fig. 2 and Ext. Figs. 1 and 2). We determined this interval as the one that minimised CoV · 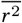, where CoV is the coefficient of variation of the yields in the experiment and 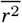 is the square of Pearson’s r of the OD and time over the interval averaged over the experiment. The latter serves to favour intervals where growth stagnates, as expected for saturation. We further imposed the following constraints on the interval: First, the interval should not begin earlier than the median of times at which growth curves in the experiment exceeded 0.7 of their preliminary yield (see above). Second, the start of the interval should be between 0.5 and 0.9 of its end time (with the beginning of the observation set to 0 s). We used a brute-force grid search for the minimisation, using 100 steps in each dimension and Powell’s method as a finisher.

We computed the growth rate by fitting a logistic growth curve within a given OD interval (horizontal grey area in Fig. 2 and Ext. Figs. 1 and 2). A simple exponential growth was included as a special case (due to a liberal constraint on the yield) and also used as a fallback if the fit of a logistic curve did not converge. We determined the interval for the fit as the one that minimised CoV · 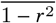, where CoV is the coefficient of variation of the growth rates in the experiment and 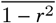 quantifies the badness of fit using the coefficient of determination r averaged over the experiment. We further imposed the following constraints on the OD interval: First, the lower end of the interval should lie between 0.001 and 0.1 of the median yield for the experiment. Second, the upper end should not exceed the yield. Third, the interval should span between 0.7 and 3 powers of ten. We minimised analogously to yield in logarithmic OD space. When using a simple exponential fit instead, the median growth variability (over both conditions) was higher and the variability reduction when supplementing trace elements was lower (data not shown).

**Ext. Fig. 1.**
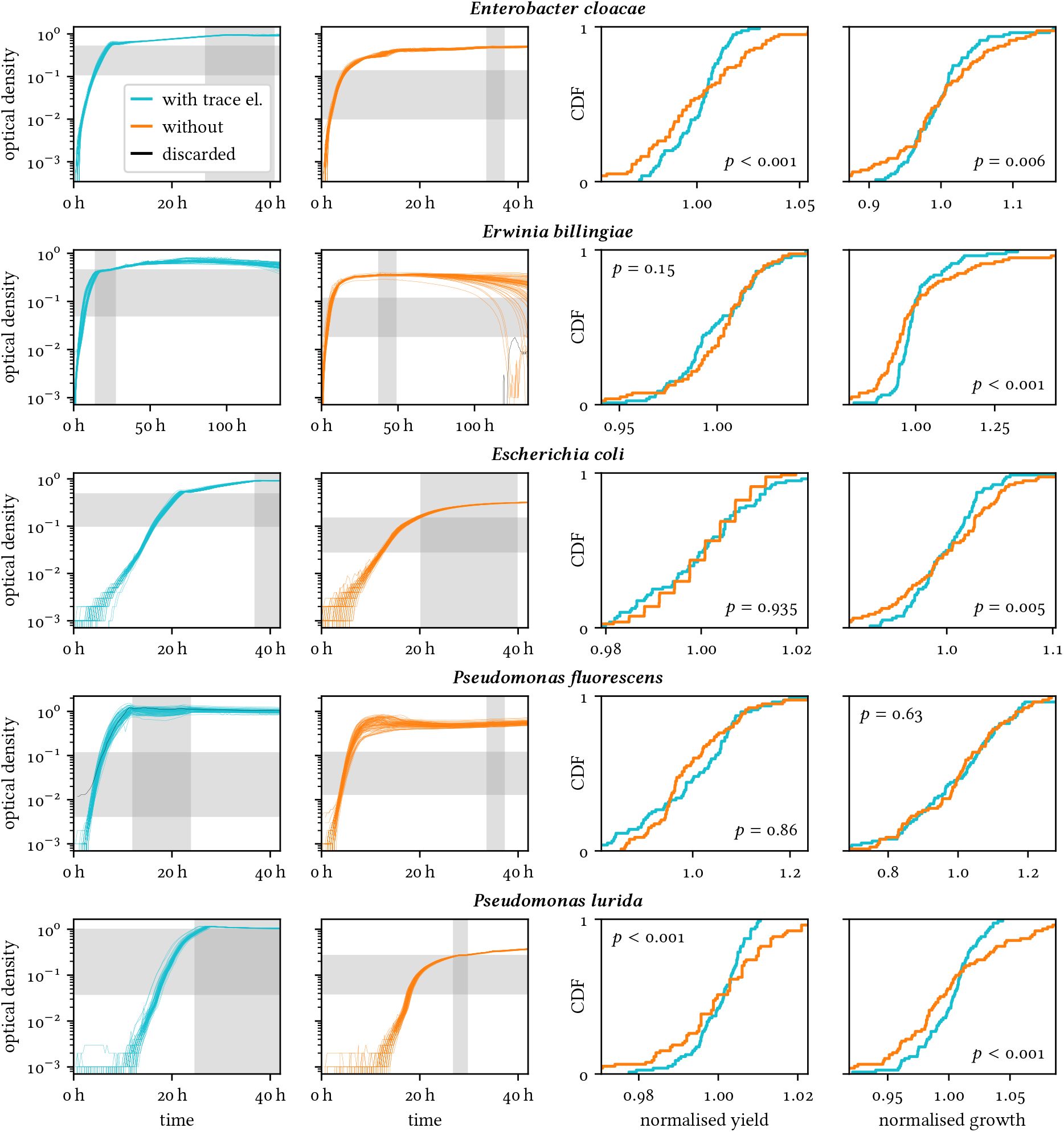
Results for all strains. Like Fig. 2 without smoothing and with discarded curves. Left: 80 growth curves recorded under identical conditions with and without trace elements for each strain. Grey areas mark the time and optical-density ranges used for determining the yield and growth rate respectively. Discarded curves (black) were not used in further analysis. Right: Distributions of normalised growth characteristics with and without trace elements with the one-sided *p* value that the coefficient of variation is lower with trace elements. Continued in Ext. Fig. 2.

**Ext. Fig. 2.**
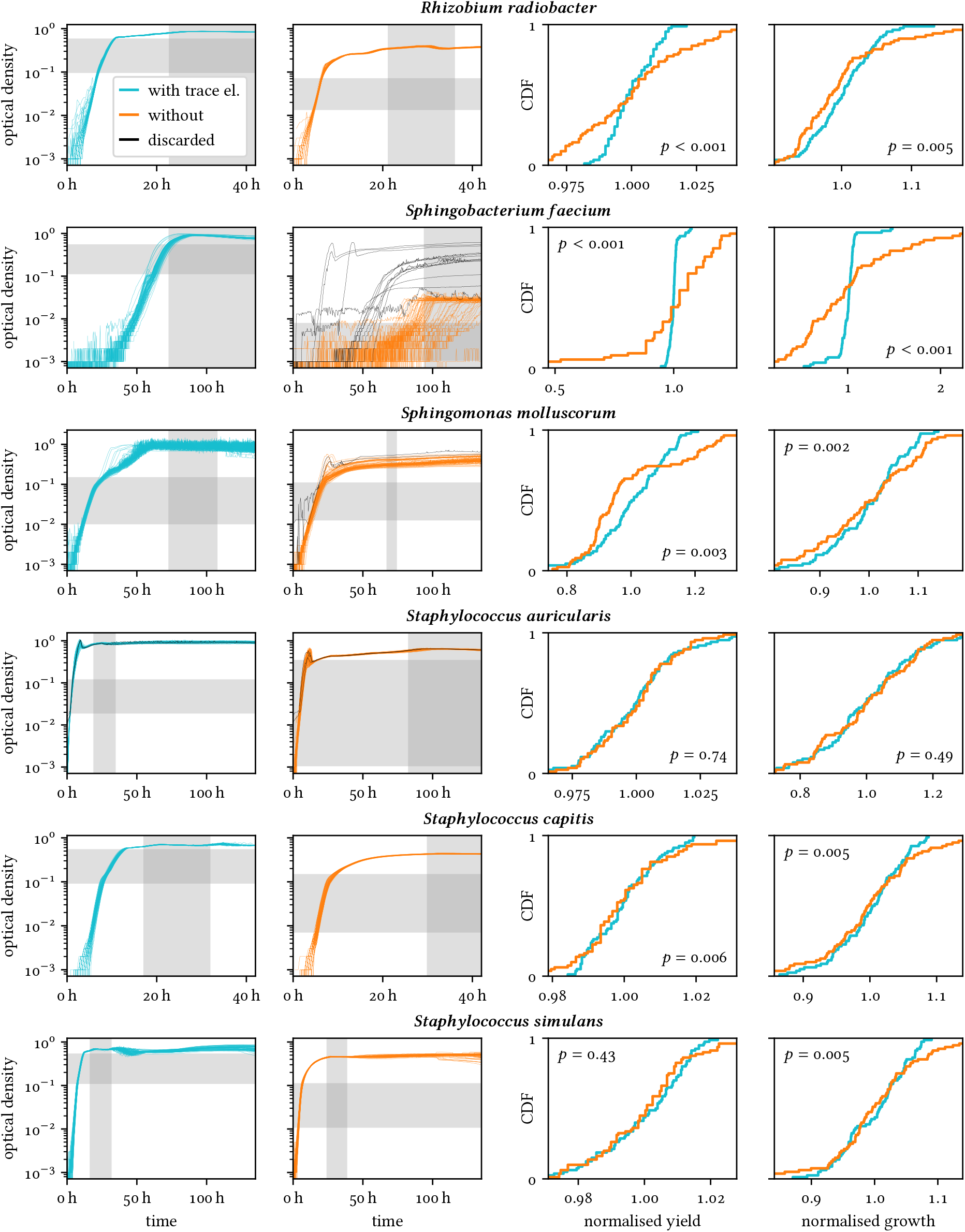
Continuation of Ext. Fig. 1.

**Ext. Fig. 3.**
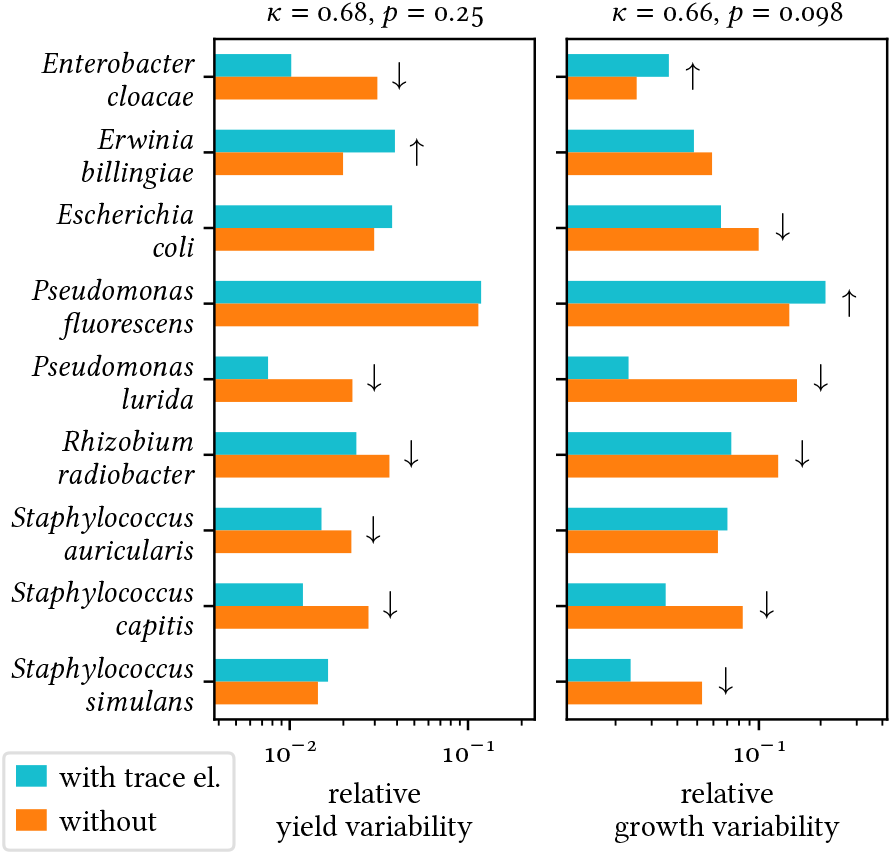
Glassware does not appear to be the cause of trace-element contamination (relevant to our experiments). Like Fig. 3, but half of the medium for each experiment was subdivided into eight bottles before being distributed over the plates (see Methods 3). If the bottles were a relevant source of contamination, we would expect an amplification of the impact of trace elements on growth variability (Fig. 3).

### 7. Summarising statistics

To statistically test whether two datasets differ in variability, we first normalised the values in each dataset by dividing by the dataset’s mean and then performed a permutation test with 10000 permutations, in which we randomly re-assigned the normalised values to the datasets (Ext. Figs. 1 and 2).

To quantify and statistically test the overall effect, we computed for each strain *i* the ratio of the coefficients of variation:

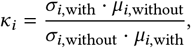

and used Wilcoxon’s signed-rank test for the null hypothesis that the median of log (*κ*_*i*_) is zero (Fig. 3). The logarithm here ensures that the symmetry conditions of the test are met. We performed this test separately for yield and growth rate to avoid pseudoreplication.

To assess a potential correlation of the variability of a yield and growth rate and their respective means (Ext. Fig. 4), we employed Pearson’s r. Since the distributions of means and variabilities do not appear to be normal, we did not use the typical test based on the *t* distribution, but instead used a permutation test with 10000 permutations, randomly re-pairing means and variabilities.

**Ext. Fig. 4.**
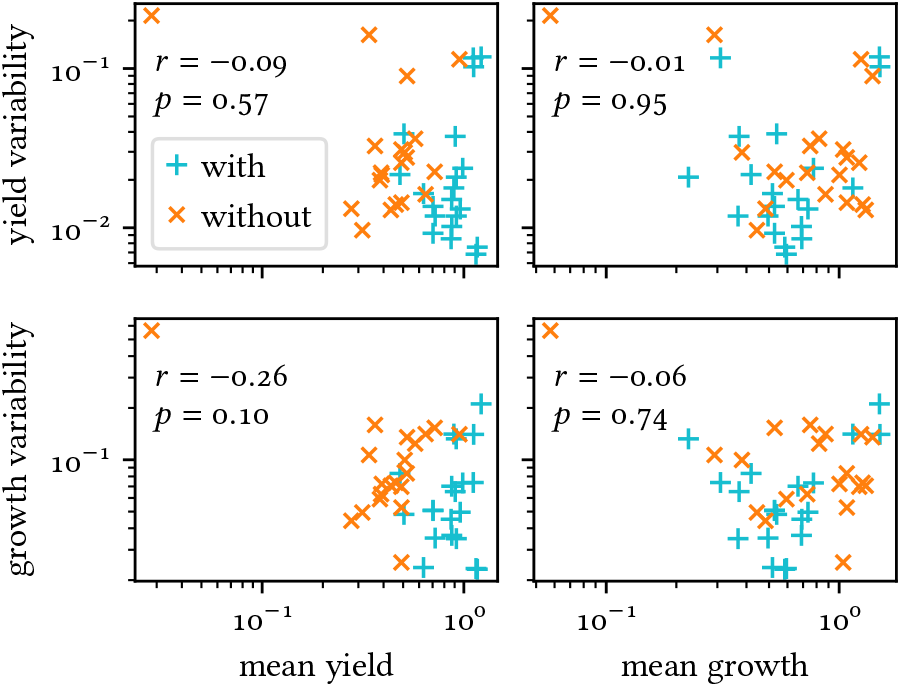
Our variability analysis is not confounded by artefacts related to the absolute value of the respective measure. Scatter plots of mean yield or growth and their variabilities for each experiment – with Pearson’s r and its two-sided *p* value (Methods 7).

### 8. Detecting trace-element contamination

To estimate the magnitude of trace-element contamination, we repeated the core measurement procedure with ultrapure water, specifically: We filled a reservoir (Thermo 1200-1300) from a water purifier (ELGA PF2XXXXM1) and transferred its contents to a microtitre plate, which we incubated and measured regularly in the spectrophotometer as described above. Afterwards, we centrifuged the contents out of the plate into the same reservoir (which we had emptied immediately after usage) and added 5% HNO_3_ (Carl Roth 2616.2) for mass spectroscopy. Finally, we poured 3 ml each into four metal-free tubes (Carl Roth, XX96.1 or VWR, 525-0629; the results showed no distinction between those). The above procedure aims at minimising the number of labware items coming in contact with all the water, as opposed to individual wells, as the latter are the most likely source of variability studied here. Note that the water did not come into contact with any glass labware. We contrasted our samples with four negative controls, for which we filled the tubes directly from the water purifier (and added 5% HNO_3_).

The elemental contents of each tube were determined via mass spectroscopy (Agilent 7700 ICP-MS) by the University of Cologne’s Biocentre MS Platform. Measurements were performed in technical triplicates (per sample), strictly following the manufacturer’s instructions using helium in the collision cell mode to minimise spectral interference. Of the trace elements from Ext. Tab. I, tungsten wasn’t measured, while results for aluminium, selenium, and molybdenum (^95^Mo and ^98^Mo) were below the detection limit for all samples. On top, other trace elements, namely lithium, vanadium, chromium, arsenic, strontium, cadmium, platinum, and lead (^206^Pb, ^207^Pb, and ^208^Pb) were measured.

**Ext. Fig. 5.**
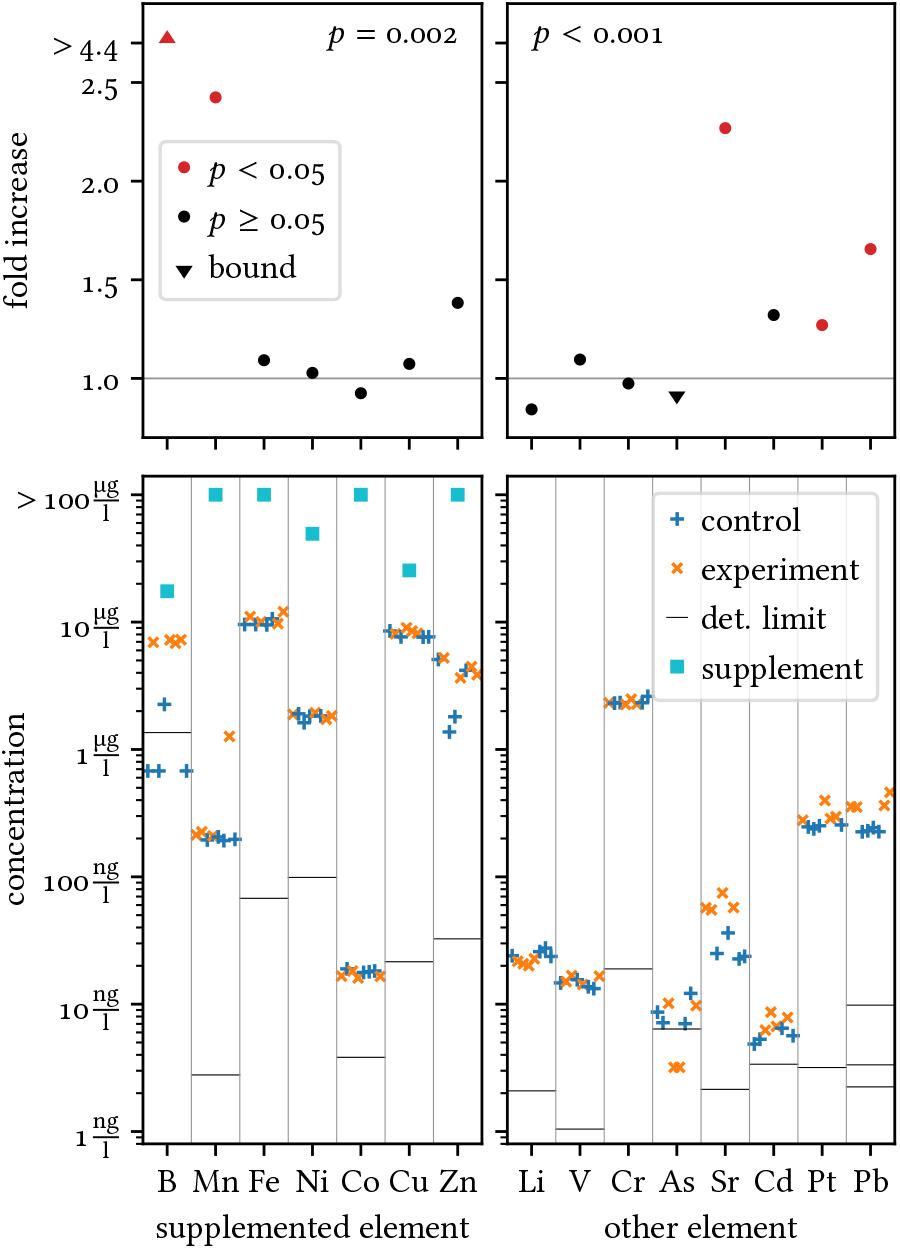
The experimental procedure or the labware leads to trace-element contamination. Left: elements contained in the employed trace-element solution (Ext. Tab. I). Right: other trace elements. Top: fold increase of trace-element concentrations in highly purified water through the experimental procedure (filling into reservoir, transferring into microtitre plates, incubating and measuring) with respect to the negative control (Methods 8). Red markers indicate that, for this particular element, the difference to the negative control is significant (*p <* 0.05, two-sided Mann– Whitney *U* test over replicates). The *p* values at the top are obtained by combining those tests over all elements (two-sided). Triangles indicate upper or lower bounds when some concentrations were under the detection limit. Bottom: absolute concentrations in the above cases (experiment and control) for each replicate, the detection limit, and when trace elements are supplied (per manufacturer’s specification, Ext. Tab. I). Values are horizontally scattered to reduce overlap. Values below the detection limit are plotted as half the detection limit.

For each element, we compared all samples with the negative controls using the Mann–Whitney *U* test (Ext. Fig. 5). To assess the overall significance, we combined the *p* values using the Mudholkar–George method^28^ – separately for investigated and additional trace elements. Results for lead were combined similarly for the three isotopes.

### 9. Software

We used routines from SciPy^29^ (Version 1.13.1) for most statistical analyses, specifically brute-force minimisation, curve fitting, Pearson’s r, and Wilcoxon’s signed-rank test. We used a Python module for combining *p* values from discrete tests^30^ (Version 1.2.3) for the tests in Methods 8 and combining them.

## DATA AND CODE AVAILABILITY

All data and analysis code for this manuscript are available in the Zenodo repository No. 12746431.

## ACKNOWLEDGMENTS

We are grateful to the entire Bollenbach group for inspiring discussions as well as B. Bagheri, C. Ernst, and T. Fink for constructive comments on previous versions of the manuscript. We thank the University of Cologne’s Biocentre MS Platform, in particular S. Ambrosius and S. Metzger for performing mass spectroscopy and help with our preparation, L. Goretzky for developing protocols for robotic automatisation, and H. Schulenburg for providing the strains from Ref. 25. This work was partly funded by DFG SFB 1310 (German Research Foundation, Collaborative Research Centre 1310, project number 325931972).

## AUTHOR CONTRIBUTIONS

G.A. and R.G. conceived and conceptualised the study. A.S.N., G.A., and R.G. designed the growth experiments with help of T.B., and A.S.N. performed them. G.A. designed and performed the contamination analysis (except mass spectroscopy). G.A. analysed the data. G.A. and T.B. coordinated and supervised the work. R.G. conducted literature research with help of G.A. G.A. wrote the manuscript with input from A.S.N., R.G., and T.B.

## Notes

https://dx.doi.org/10.5281/zenodo.12746431

### Competing Interest Statement

The authors have declared no competing interest.

### Summary of Updates

Better explanation of the mechanistic backbone, distribution of effect sizes, and relevance of the investigated phenomenon.

